# TCR_Explore: a novel webtool for T cell receptor repertoire analysis

**DOI:** 10.1101/2022.11.03.514642

**Authors:** Kerry A. Mullan, Justin B. Zhang, Claerwen M. Jones, Shawn J. R. Goh, Jerico Revote, Patricia T. Illing, Anthony W. Purcell, Nicole L. La Gruta, Chen Li, Nicole A. Mifsud

**Author notes:** To whom correspondence should be addressed: Kerry A. Mullan, Nicole A. Mifsud. **Biographical notes**. **Kerry A. Mullan** is a Ph.D. candidate in the Department of Biochemistry and Molecular Biology, Monash University. Her research focuses on Type IV hypersensitivity drug (*e.g*. carbamazepine) reaction that explored genomic/transcriptomic markers, functional T cell work, as well as bioinformatics program development. **Justin B. Zhang** is a Ph.D. candidate in the Department of Biochemistry and Molecular Biology, Monash University. His research focuses on how the CD4 and CD8 co-receptors regulate T cell receptor recognition, instruct T cell development and shape T cell receptor repertoires. **Claerwen M. Jones** is a post-doctoral research fellow at the Biomedicine Discovery Institute and Department of Biochemistry & Molecular Biology, Monash University. Her current research focuses on T cell responses in autoimmune diseases. **Shawn J. R. Goh** is a Ph.D. candidate in the Department of Biochemistry and Molecular Biology, Monash University. His research focuses on Type IV drug hypersensitivity (e.g. anticonvulsants, antibiotics) that explored how penicillins alter the immunopeptidome of T cells and the repertoire of TCRs involved in such reactions. **Jerico Revote** is a Research DevOps Systems Engineer at the Monash eResearch Centre (MeRC), Monash University with a background in cloud infrastructure and solution design. He is the technical lead in the deployment and operations of the Monash Secure eResearch Platform (SeRP). He is also the activity lead for the digital cooperatives capability of the Monash eResearch Centre. He has established a wide range of technology research capabilities used by hundreds of Australian and international researchers to date. **Patricia T. Illing** is a Group Leader in the Biomedicine Discovery Institute and Department of Biochemistry & Molecular Biology, Monash University, Australia. Her research centres on understanding peptide and small molecule presentation by MHC, and MHC-like, molecules in the context of infection, cancer and adverse drug reactions. **Anthony W. Purcell** is a Professor and head of immunopeptidomics and quantitative proteomics in the department of Biochemistry and Molecular Biology at Monash University. His current research focuses on using cutting edge proteomics to address a wide variety of immune questions related to human immunology, molecular virology, structural and functional immunology. **Nicole L. La Gruta** is a Professor and head of the T cell development and function laboratory in the Department of Biochemistry and Molecular Biology and the Immunity Program in the Biomedicine Discovery Institute at Monash University. Her current research focuses on understanding the molecular bases of healthy and dysfunctional (ageing, autoimmunity) T cell responses. **Chen Li** is a Research Fellow in the Biomedicine Discovery Institute and Department of Biochemistry and Molecular Biology, Monash University. His research interests include systems proteomics, immunopeptidomics, personalized medicine, and experimental bioinformatics. **Nicole A. Mifsud** is Group Leader in the Biomedicine Discovery Institute and Department of Biochemistry & Molecular Biology, Monash University, Australia. Her current research program explores T cell-mediated immune responses in transplant rejection, as well their contribution in human disease settings such as viral infections, adverse drug reactions and cancer.

## Abstract

T cells expressing either alpha-beta or gamma-delta T cell receptors (TCR) are critical sentinels of the adaptive immune system, with receptor diversity being essential for protective immunity against a broad array of pathogens and agents. Programs available to profile TCR clonotypic signatures can be limiting for users with no coding expertise. Current analytical pipelines can be inefficient due to manual processing steps, open to data transcription errors and have multiple analytical tools with unique inputs that require coding expertise. Here we present a bespoke webtool designed for users irrespective of coding expertise, coined ‘TCR_Explore’, incorporating automated quality control steps that generates a single output file for creation of flexible and publication ready figures. TCR_Explore will elevate a user’s capacity to undertake in-depth TCR repertoire analysis of both new and pre-existing datasets for identification of T cell clonotypes associated with health and disease. The web application is located at https://tcr-explore.erc.monash.edu for users to interactively explore TCR repertoire datasets.

**Key Points:**

- Bespoke program for non-specialists in computerised methodologies for deep exploration of TCR repertoire analysis
- Automated QC and analysis pipelines for Sanger based TCR sequencing coupled with immunophenotyping, with the capacity for integration of other sequencing platform outputs
- Automated summary processes to aid data visualisation and generation of publication-ready graphical displays

## Introduction

Conventional alpha-beta (αβ) T cells and unconventional gamma-delta (γδ) are critical sentinels of the immune system that are equipped with a molecular armory to detect, engage and eliminate abnormal cells[1–3]. Both αβTCR and γδTCR are heterodimeric protein containing an α- and a β-chain or a γ- and a δ-chain, respectively. The α- and γ-chain, termed *TRA* or *TRG*, are encoded by one variable (*V*), one joining (*J*) and one constant (*C*) gene, whilst the β- and δ-chain, termed *TRB* or *TRD*, is encoded by one *V*, variable diversity (*D*) genes (up to three), one *J* and a *C* gene [4, 5]. Recombination of various *V(D)J* genes, including incorporation of non-template, results in distinct TCR complementarity determining region (CDR) 3 protein sequences[4, 6], and leads to highly diverse TCR repertoire (*e*.*g*. 10^6^ to 10^8^ and unique functional αβTCR clonotypes[7]). Determining the composition and diversity of the TCR repertoire associated with different human diseases is an important step in rationalised clinical interventions or development of T cell-based immunotherapies. For example, we have applied single-cell αβTCR[8] and γδTCR[9] gene analysis to decipher MHC-restricted TCR signatures associated with herpesvirus infection[10, 11], autoimmune diseases such as rheumatoid arthritis[12], heterologous immunity[13] and drug hypersensitivity (αβTCR[14, 15]). Whilst the acquisition pipeline of single-cell TCR data by Sanger sequencing is relatively standardised (**Figure 1**), the downstream quality control (QC) processes including filtering poor quality sequences and manual pairing of αβ or γδ TCR chains, as well as verification to ensure data transcription accuracy remain labour intensive. Similarly, single chain data aligned utilising specific software or paid services (*e*.*g*. ImmunoSEQ[16], MiXCR[17]) can also require post-alignment QC processes that include filtering out poor quality sequences. Moreover, visualisation of TCR repertoire data often requires additional manual reformatting steps to conform to input requirements of figure generation pipelines such as the Circos® online tool[18], downloadable coding-based programs (*e*.*g*. TCRdist, VDJtools)[17, 19–25], or subscribed non-TCR specific statistical programs (*e*.*g*. GraphPad Prism 9 [GraphPad, Software, San Diego, CA, USA] or Microsoft®Excel® [Microsoft, Redmond, WA, USA]). Often these applications are restricted to either a single TCR chain analysis[20, 23] or paired TCR chain analysis[19]. Hence, there is an unmet need for an application that requires minimal to no coding expertise, automation of manual processes, elimination of data transcription errors, as well as improved flexibility of data analysis and figure generation.

**Figure 1.**
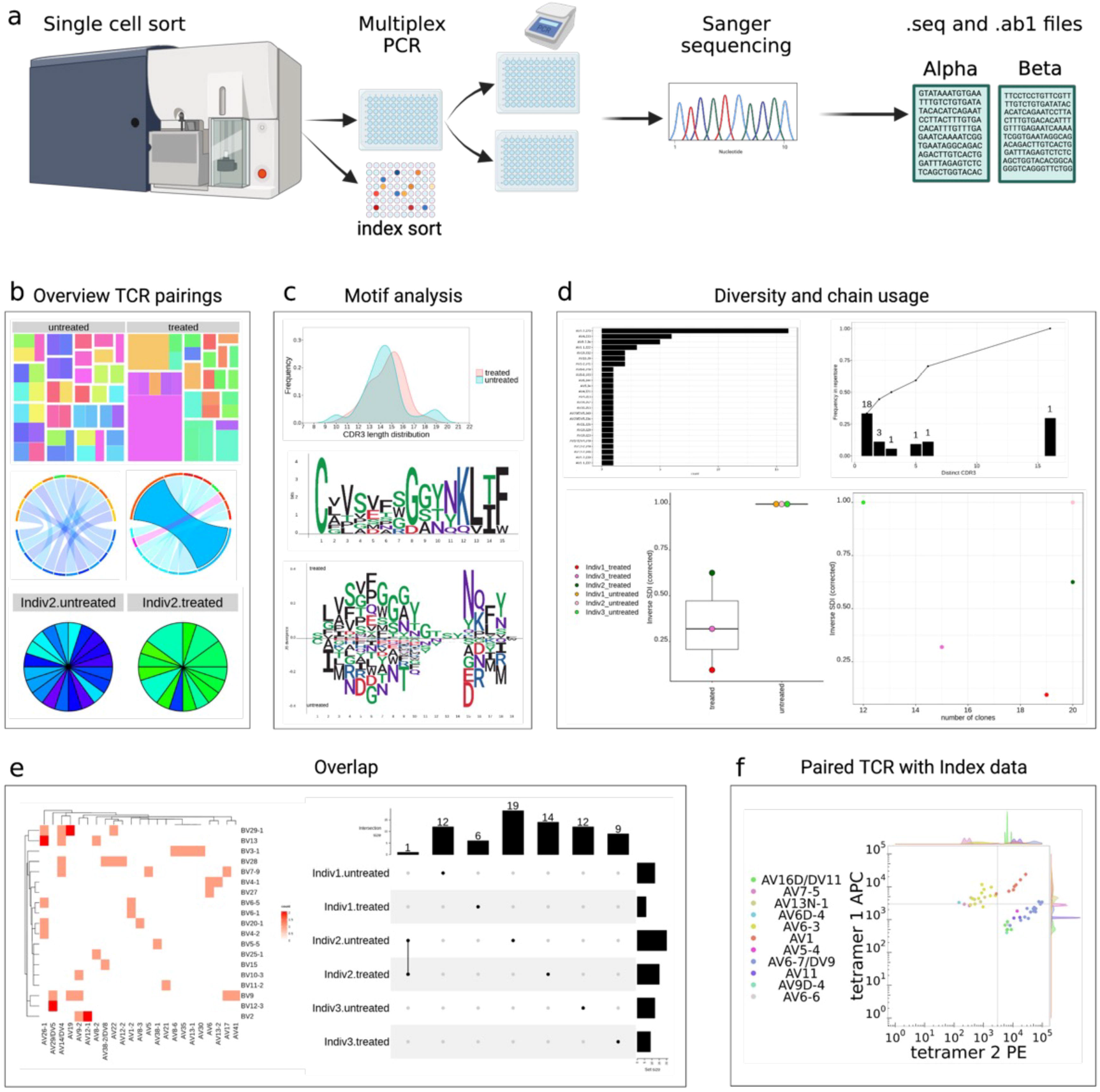
Traditional Sanger sequencing pipeline. **(A)** Targeted T cells are single-cell sorted into 96 well plates by flow cytometry. Single cells undergo reverse transcription to cDNA that is then used as the template for amplification of selected *TCR* genes by multiplex nested PCR. Following a PCR clean-up step and fluorescent labelling of dNTPs, the amplified DNA undergoes Sanger sequencing, which produces two output files (.seq and .ab1). (**B**) ‘Overview of TCR pairing’ panel; (**top**) treemap, (**middle**) chord diagram, (**bottom**) pie chart. (**C**) ‘Motif analysis’ panel; (**top**) length distribution, (**middle**), single length motif plot, (**bottom**) aligned motif plot. (**D**) ‘Diversity and chain usage’ panel; (**top left**) chain usage, (**top right**) frequency of each clonotype, (**bottom left**) inverse Simpson diversity index (SDI), (**bottom right**) total number of clones. (**E**) ‘Overlap’ group comparison panel; (**left**) heatmap or (**right**) upset plot. (**F**) ‘Paired TCR with Index data’ panel; dot plot of the functional TCR sequence and two immunophenotyping markers. Figure created using BioRender (BioRender.com).

Here we present TCR_Explore, a Shiny R application available on an open-access webserver (http://tcr-explore.erc.monash.edu) that analyses and visualises TCR repertoire data. TCR_Explore introduces workflows using an automated process that pairing of αβ or γδ TCR chains from Sanger sequencing pipelines and facilitates interrogation of linked flow cytometric index data for immunophenotyping analyses. Additionally, TCR_Explore facilitates conversion and filtering of non-Sanger alignment pipelines (*e*.*g*. ImmunoSEQ, MiXCR) to the required TCR_Explore format. Moreover, an automated summarisation process from a single input file enables the visualisation of complex data sets and creation of a variety of publication ready figures. Thus, TCR_Explore is a powerful platform for routine analysis of TCR repertoire data of new and reanalysis of pre-existing datasets for greater insight by users irrespective of coding expertise.

## Methods

### Data and code availability

The demonstration data is from Mifsud *et al*. (2021)[14] and Lim *et al*. (2021)[12]. The local version of ‘TCR_Explore’ and all the raw data files and processed datasheets are located on GitHub https://github.com/KerryAM-R/TCR_Explore in the test-data section.

### Single cell sorting and multiplex nested PCR for amplification of TCR chain genes

TCR_Explore was developed for the QC and analysis of Sanger sequencing data generated following multiplex nested PCR (**Figure 1A**). Briefly, this included a single cell sort, with and without FACSort index data, into position A1 to H10 of a 96 well plate. Followed by multiplexed PCR of either the TCRα and TCRβ or TCRγ and TCRδ[9, 10, 12, 14]. Ideally, sample labelling should follow IndividualID.groupChain-initialwell (*e*.*g*. T00020.IFNB-A1). This naming can be either added in the .seq to .fasta conversion step or prior to the pairing process. Sanger sequencing generates two outputs files, .seq and .ab1, that contains the sequencing and chromatogram information, respectively.

### Quality Control

#### Step 1: Alignment of TCR chain sequences using IMGT

Sanger sequencing .seq output files need to be converted into a .fasta file, which can be performed using TCR_Explore (recommend 50 sequences per file) (**Supplementary data 1**). The .fasta file is then uploaded onto the international ImMunoGeneTics information system® (IMGT)[4, 26] website (https://www.imgt.org/IMGT_vquest/input), which aligns a maximum of 50 sequences at a time. To download the Vquest.xls file, the user selects the relevant species (*e*.*g*. Homo sapiens), receptor type or locus (*e*.*g*. TR), followed by section “C. Excel file” containing only the ‘Summary (1)’ and ‘Junction (6)’ tabs.

#### Step 2: TCR_Explore Quality control

##### Creating QC file

Upload the Vquest.xls file into the ‘QC→IMGT (Sanger Sequencing)’ tab, select the dataset ‘own_data’ and ‘Select file for IMGT datafile’ using the browse function to upload a file. This will create a downloadable IMGT_onlyQC.date.csv file that contains necessary information for either the TCR_Explore QC process or for compatible files for use in external programs.

##### Chromatogram quality and sequence functionality

The program adds columns required for the QC process including, ‘V.sequence.quality.check’ (column Q), ‘clone_quality’ (column R) and ‘comments’ (column S) (**Supplementary Table 1**). The ‘V.sequence.quality.check’ (Column Q) flags IMGT outputs that were not aligned, unproductive in ‘V-DOMAIN Functionality’ (Column C), if there were either V or J identity issue (<90% identity) from ‘V-REGION identity %’ (column E) and ‘J-REGION identity %’ (column G), respectively or ‘No issue flagged by IMGT’. Based on the test data[14], 151 sequences reported ‘No issue flagged by IMGT’, had high quality chromatograms with productive sequences, and therefore were designated as a ‘pass’ in the ‘clone_quality’ (column R), while the 136 remaining sequences were designated as ‘NA’. The user manually interrogates the chromatogram of each .ab1 files using either TCR_Explore or external programs such as Chromas (Windows; http://technelysium.com.au/wp/chromas/), FinchTV (Mac/Windows; https://digitalworldbiology.com/FinchTV) or another chromatogram visualisation software. The user needs to fill in the ‘comments’ column on if the sequence is either of poor quality (*e*.*g*. T00024.IFNA-A3_A07.ab1 contained two sequences) or high quality (*e*.*g*. E10630.CD8A-A5_B01.ab1) (**Supplementary Table 1**). All sequences that are low quality need to be designated a ‘fail’ in the ‘clone_quality’ (column R). Next the user needs to replace the remaining ‘NA’ as either ‘pass’ or ‘fail’ depending on whether the clone is productive or not (*i*.*e*. stop codons or frameshift), and If these sequences can be manually resolved (*e*.*g*. insert missing base pair in low quality regions). The user can create a new file .fasta file with the resolved sequence(s) with ‘man’ added to the end of header (*e*.*g*. >T00020.IFNA**-**A1_A1.seq#1man), as this will not impact the pairing process. The original sequences will ‘fail’, while the manually altered sequences will be designated a ‘pass’. This ensures that all alterations are documented and traceable. Overall, the ‘clone_quality’ will be filled in by the user as either pass (*e*.*g*. productive, in-frame, no stop codons) or fail (non-productive, out-of-frame, stop codons, no rearrangement, two sequences, no result) based on the sequence information. The user can document the ‘fail’ reason in the ‘comments’ column S. This process is repeated for all Vquest.xls sequence files, and the data is combined into a single .csv file for downstream TCR chain pairing. A video example of the quality control process is located in ‘Tutorials →Quality control information (includes video tutorial)’.

#### Step 3: Creating the paired TCR file

Upload the completed QC.csv file into the ‘QC → Paired chain file’ tab, select the dataset ‘own_data’ and ‘Completed QC file (.csv)’ using the browse function to upload a file to create the paired_TCR.csv file (**Supplementary Table 2**). The user can select either alpha-beta or gamma-delta chains as well as the Information included (*e*.*g*. Summary+JUNCTION). Only paired chains that have a ‘pass’ assigned will included in the final functional paired TCR repertoire file. The pairing process is based on the IndividualID.groupChain-initialwell. Additionally, the T cell receptor (TR) abbreviation was removed (*e*.*g*. TRAV = AV). Moreover, the program adds several columns to the end of the file that shows the genes without the allele for AJ, AV, AVJ, BJ, BV, BD, BVJ, BVDJ, AVJ.BVJ, and AVJ.BVDJ or the γδTCR equivalent (**Supplementary Table 2**). This will create a downloadable .csv file (*e*.*g*. paired_TCR.csv) that contains necessary information for both the ‘TCR analysis’ and ‘Paired TCR with index data’ sections. There is also an option to download the cleaned ‘single chain file’ if pairing is not needed or the Tab space variable (TSV) or .tsv file for use in TCRdist[19].

### Conversion of alternate TCR data outputs to TCR_Explore format

TCR_Explore can convert TCR repertoire data from other alignment programs into a compatible format. For ImmunoSEQ® we utilised data from Heikkila *et al*. (2021)[27], this process rearranges the file so that the count data is in column A (renamed as cloneCount), keeps the in-frame sequences only, removes empty columns and missing information from either the V or J genes. For MiXCR[17], the program removes sequences with stop codons or frameshifts. For sequencing data not aligned through either ImmunoSEQ® or MiXCR, use the ‘Other’ in the ‘Input type’ dropdown menu, as this contains the generic filtering and conversion functions. A video example of the functionality is provided in ‘QC → Convert to TCR_Explore file format → Video of the conversion process’.

### TCR analysis

The ‘TCR analysis’ tab includes features to aid in figure generation and summary statistics. Users can select either test αβTCR data[14] denoted as “ab-test-data2” or they can upload the single or paired TCR .csv file or from another QC processed TCR dataset (*e*.*g*. MiXCR[17]). Interrogation of TCR data is achieved via four distinct analysis platforms (Overview of the TCR, Motif analysis, Diversity and chain usage, Overlap). Each section is further subdivided using tabs or dropdown menus to alter graphing parameters and displays for each figure. In total, there are 14 distinct figures that can be generated in ‘TCR analysis’ section and one in the ‘Paired TCR with Index data’ section.

#### (i) Overview of TCR pairing

‘Overview of TCR pairing’ tab includes a downloadable summary table for TCRdist3[24], and three analytical graphs: treemap, chord diagram and pie chart. There are common automated and customisable features of the plots which include: ordering the groups, customisable colours, font type, drop-down menus to change the desired comparison. The drop-down menus enable the user to quickly change their comparison from single chain analysis (*e*.*g*. TRAV vs TRAJ) to paired chain analysis (*e*.*g*. TRAV-TRAJ vs TRBV-TRBJ) without the need to manually alter the file. Additionally, the chord diagram included options for selective labelling (*e*.*g*. Label or colour selected clone/s). A video example of the functionality is available in ‘Tutorials → Video examples → Overview of TCR pairing’.

#### (ii) Motif Analysis

There are four sub-tabs in the ‘Motif analysis’ section which presents motif plots based on the unique CDR3 sequences. The first tab is the ‘CDR3 length distribution’, which includes a histogram that can be colour coded by a specific column (*e*.*g*. AVJ) or as a density plot that shows the overlap of the groups. The three other sub-tabs show either the ‘Motif (amino acid)’ and ‘Motif (nucleotide sequence)’ for single lengths, while the ‘Motif (AA or NT alignment)’ uses muscle[28] to align the sequences. Both the ‘Motif (amino acid)’ and ‘Motif (AA or NT alignment)’ can compare the differences of two motifs using subtractive analysis. Like the online version of muscle, we restricted sequence alignment to 500 sequences to prevent server timeout issues. A video example of the functionality is provided in ‘Tutorials → Video examples → Motif analysis’.

#### (iii) Diversity and chain usage

There are two tabs in the ‘Diversity and chain usage’ section. The first tab ‘Chain bar graph’ has three distinct graphs of either the chains used per group, the frequency of the repertoire per group and a stacked bar graph. For the frequency graph, the x-axis represents the number of times a clone was observed, the numbers above the bars represent the unique clones, and the line represents the cumulative frequency. The second tab is the ‘Inverse Simpson Index’, which is used to calculate the changes in diversity. The larger the Inverse Simpson Index, the more diversity. There are two graphs available; (1) shows the index vs the selected group and (2) showcases the index vs either the number of total clones or unique clones. Graph (2) is used to check the total number of sequences is not biasing the results. If two groups are being compared, a standard t-test calculation is available. For more complex statistical methods, such as ANOVA, a third-party program is required, therefore the Inverse Simpson index table is downloadable for this purpose. Further interrogation of the data can also be conducted using TCRdist[19] or TCRdist3[24]. A video example of the functionality is in ‘Tutorials → Video examples →Diversity and chain usage’.

#### (iv) Overlap

The ‘Overlap’ section enables users to compare multiple groups using either a heatmap or upset plot. The heatmap compares chain usage from either single or multiple individuals, whilst an upset plot can display the overlap of up to 31 groups, which is a restriction of the package used[29]. These comparisons highlight whether the TCR repertoires are of a public or private nature. The upset plot table data is also downloadable. A video example of this functionality is in ‘Tutorials → Video examples → Overlap’.

### Paired TCR with Index data

This three-step process is showcased as a video in ‘Tutorials → Video examples → Paired TCR with Index data’.

#### Step 1. Merging the paired TCR with Index data (QC process 1)

The ‘Paired TCR with Index data’ section automates the merging of the paired clone file with the corresponding .fcs file. A background file converts the .fcs xloc and yloc (*e*.*g*. 0,0) values to A1 to H10 values to enable merging. This process is limited to one plate at a time as only one .fcs can be uploaded. However, there is no need to reformate the QC paired TCR .csv file. The user needs to select the group, individual and if there were multiple plates. The user can then copy all the samples into one .csv file.

#### Step 2. Data cleaning steps (QC process 2)

The next quality control step occurs in ‘Data cleaning steps’ to covert the negative fluorochrome values to small positive values required for log transformation, and was the method utilised in Lim *et al*. (2021)[12]. To create the colour scheme for the file, the users can select all necessary columns, but must not select fluorochrome columns, well, cloneCount columns as it will not summarise. After downloading, the user can alter names of the fluorochromes, which is restricted to alphabetic characters and numbers.

#### Step 3. Generation of the analytical plot

Next, in the ‘TCR with index data plot’, the user uploaded the cleaned file from step 2. The user can select any of the fluorochromes to display on the graph. There are over 20 customisable features including the size, colour, and shape of each dot as well as text size and font. These features are either located in the side panel or above the plot, so the user can readily visualise all changes. The figure can be downloaded as either a PNG or PDF.

### R packages

TCR_Explore is an R-based Shiny application constructed using various R packages including: “tidyverse” (version 1.3.1)[30], “ggplot2” (version 3.3.5)[31], “ggrepel”(version 0.9.1)[32], “shiny” (version 1.7.1)[33], “shinyBS” (version 0.61)[34], “gridExtra”(version 2.3)[35], “DT”(version 0.20)[36], “plyr” (version 1.8.6)[37], “dplyr” (version 1.0.7)[38], “reshape2” (version 1.4.4)[39], “treemapify” (version 2.5.5)[40], “circlize” (version 0.4.13)[41], “motifStack” (version 1.36.1)[42], “scales” (version 1.1.1)[43], “flowCore” (version 2.4.0)[44], “readxl” (version 1.3.1)[45], “RcolorBrewer” (version 1.1-2)[46], “randomcoloR” (version 1.1.0.1)[47], “colourpicker” (version 1.1.1)[48], “ComplexHeatmap” (version 2.9.4)[29], “muscle” (version 3.34.0)[28], “DiffLogo” (version 2.16.0)[49], “vegan” (version 2.5-7)[50], “VLF” (version 1.0)[51], “ShinyWidgets” (version 0.7.0)[52], “showtext” (version 0.9-5)[53], “ggseqlogo” (version 0.1)[54], “markdown” (version 1.1)[55], “rmarkdown” (Version 2.14)[56], and “sangerseqR” (Version 1.32.0)[57]. All dependent packages used to run the program are in ‘Tutorials → Session info’.

## Results

### Interrogation of TCR repertoires

TCR_Explore is a web-application for TCR repertoire quality control and analysis. The automated QC process was created to aid cleaning and pairing of αβTCRs or γδTCRs from different sequencing alignment pipelines (**Figure 1A**) to create a single input file for TCR_Explore. Importantly, both single chain and paired chain TCR repertoire can be interrogated.

The ‘TCR analysis’ tab has four subsections for repertoire analysis. The first section enables data visualisation of TCR repertoire profiles via treemaps, chord diagrams or pie charts (**Figure 1B**) and generates a summary table. The second section evaluates CDR3 length distributions and amino acid motifs (both single length and consensus sequences) can be plotted for comparative analyses (**Figure 1C**). The third section examines changes in repertoire diversity and chain usage via inverse Simpson diversity index (SDI) values or frequency plots (**Figure 1D**). The fourth section facilitates a comparative group overlap analysis using heatmaps or upset plots (**Figure 1E**), with a downloadable table output. Collectively, TCR_Explore has enhanced flexibility to perform TCR repertoire analysis and aid in directing the next stages of analysis or experimentation.

### Showcasing TCR immunophenotypes

Unlike existing tools, TCR_Explore enables merging of functional TCR repertoire data with phenotypic expression, which is critical for validation of T cell biomarkers[12]. Previously, manual merging of paired TCR repertoire data with phenotypic markers (*i*.*e*. index data) collected during the FACSort was cumbersome and prone to data transcription errors (**Table 1**). In addition, manual conversion of negative expression values for log transformation was also required, as well as the need to select the phenotypic comparison before generating a biexponential figure in GraphPad®[12] (**Table 1**). Due to the time-consuming nature of this workflow, few studies have utilised this validation process[12, 58]. Here, the ‘Paired TCR with Index data’ tab automates these processes to generate a biexponential plot (**Figure 1F**). Drop-down menus improve flexibility for display of selected phenotypic markers, providing a detailed assessment of T cell specific biomarkers.

### Improved quality control process for pairing TCR chain genes

TCR_Explore, utilising the R statistical language, automates numerous manual QC processes as depicted in **Table 1**. This process includes pairing the separate TCRα and TCRβ or TCRγ and TCRδ chains, produced from Sanger sequencing output files (.seq). Importantly, our program also includes automated merging the functional paired TCR file with corresponding phenotype index data (.fcs) file (**Table 1**; **Supplementary Table 3**). Both QC merging process, based on the naming convention (IndividualID.groupChain-initialwell), thereby reduce the time needed to create an analysis file, with substantial reduction of data transcription errors in the QC process.

### Automated summarisation reduces errors and enables flexible figure generation

Previous workflows for TCR repertoire analysis and visualisation involved the use of multiple tools, each requiring specific file formats[17–25]. This process is time intensive, vulnerable to data transcription errors, and inflexible with respect to incorporating data updates or changes to the comparison of interest (*e*.*g*. TRAV vs TRAJ to TRAV vs TRBV). Additionally, these programs include limitations in plot customisation (*i*.*e*. font choices or colouring of specific chains)[18, 20], inflexible in their export functions (*e*.*g*. inability to specify height/width; PNG or PDF only)[18, 20] and restricted to single chain analysis[20, 21]. Therefore, there is an unmet need to develop a program to overcome these limitations.

To remove these reformatting processes and reduce errors, TCR_Explore was designed to rely on a single integrated input file for the generation of all data figures (**Table 2**). The program promptly summarises the data based on researcher inputs located in the Shiny R interface, thereby eliminating the need for creating multiple files and removal of data transcription errors. The input file structure and user-friendly interface allows flexibility, particularly if samples are added or removed. Overall, TCR_Explore has improved flexibility and breadth of analysis, thereby enhancing opportunity for TCR repertoire discoveries.

### Interface with existing programs for extended data analysis

One statistical program that uses paired TCR data is TCRdist[19], which has many features including identifying the origin of TCR sequences, principal component analysis, αβTCR epitope-specific repertoires, and a robust TCR diversity statistic. These features were beyond the scope of TCR_Explore, therefore we included an interface that serves to provide compatible outputs for TCRdist via the generation of a TSV file (**Supplementary data 2**). Additionally, we included a .csv output that was compatible with further analysis using TCRdist3[24] (**Supplementary data 3**). Moreover, the TCR data outputs from other programs and pipelines including: iRepertoire (deep-sequencing of TCR repertoire[59]), ImmunoSEQ[16], and MiXCR[17]. These external TCR data, once converted (see ‘QC → Convert to TCR_Explore file format’), can be imported into the TCR_Explore ‘TCR analysis’. Overall, TCR_Explore is compatible with external programs and data pipelines.

### Extended analysis of drug-induced human αβTCRs reveals a central residue for TCR activation

In our recent study[14], we examined the carbamazepine-induced αβTCR profile of patients who had previously experienced either Stevens-Johnson syndrome or toxic epidermal necrolysis following prescription of an anti-seizure medication, carbamazepine. These severe allergic responses are classified as T cell-mediated drug hypersensitivity reactions that principally target the skin following carbamazepine exposure via TCR activation[59, 60]. To profile the TCR repertoire of our patient cohort, peripheral blood mononuclear cells (PBMC) were *in vitro* stimulated for 14 days with 25 μg/mL carbamazepine. On day 14, drug-induced T cells were restimulated with HLA allotype matched antigen presenting cells (APC) in the absence or presence of carbamazepine, and single-cell sorted based on the production of the proinflammatory cytokine IFNγ into two subsets: (i) CD8^+^IFNγ^+^ (IFN activated group) or (ii) CD8^+^IFNγ^-^ (CD8 non-activated group). In our initial publication, we reported the drug-induced TCR clonotypes were highly focused to a few TCR clonotypes and this TCR usage was private amongst the cohort. Using TCR_Explore, reanalysis of the pre-existing dataset demonstrated a capacity to not only replicate the published findings but also to interrogate the data to new depths and reveal nuances not previously appreciated.

For the reanalysis, we examined three patients E100630, T00016 and T00024. Firstly, for E10630 drug-specific TCRs (TRAVJ-TRBVJ) were visualised using chord diagrams in TCR_Explore to demonstrate recapitulation of the Circos® plots shown in the publication[14] (**Figure 2a**). Next, we confirmed via an upset plot that the paired αβTCR chains were specific to the individual patient (*i*.*e*. private TCR repertoire) by comparing multiple individuals TCR repertoires (**Figure 2b**). TCR_Explore also enabled greater flexibility for data display using either treemaps (**Figure 2c**) or pie charts (**Figure 2d**), which represent alternatives to chord diagrams. Additionally, examination of diversity based on unique sequences showed a decrease in TCR clonotypes in the IFN activated group compared to the CD8 non-activated group. This reduced diversity was represented by a significant reduction of the sample size corrected inverse SDI score (p=0.026; paired t-test; one-tailed), which highlighted that all three individuals expressed carbamazepine-induced TCR clonotypes (**Figure 2e**). TCR_Explore automation provided the opportunity for novel findings including identification of additional clonal TCR sequences that warrant further functional validation.

**Figure 2.**
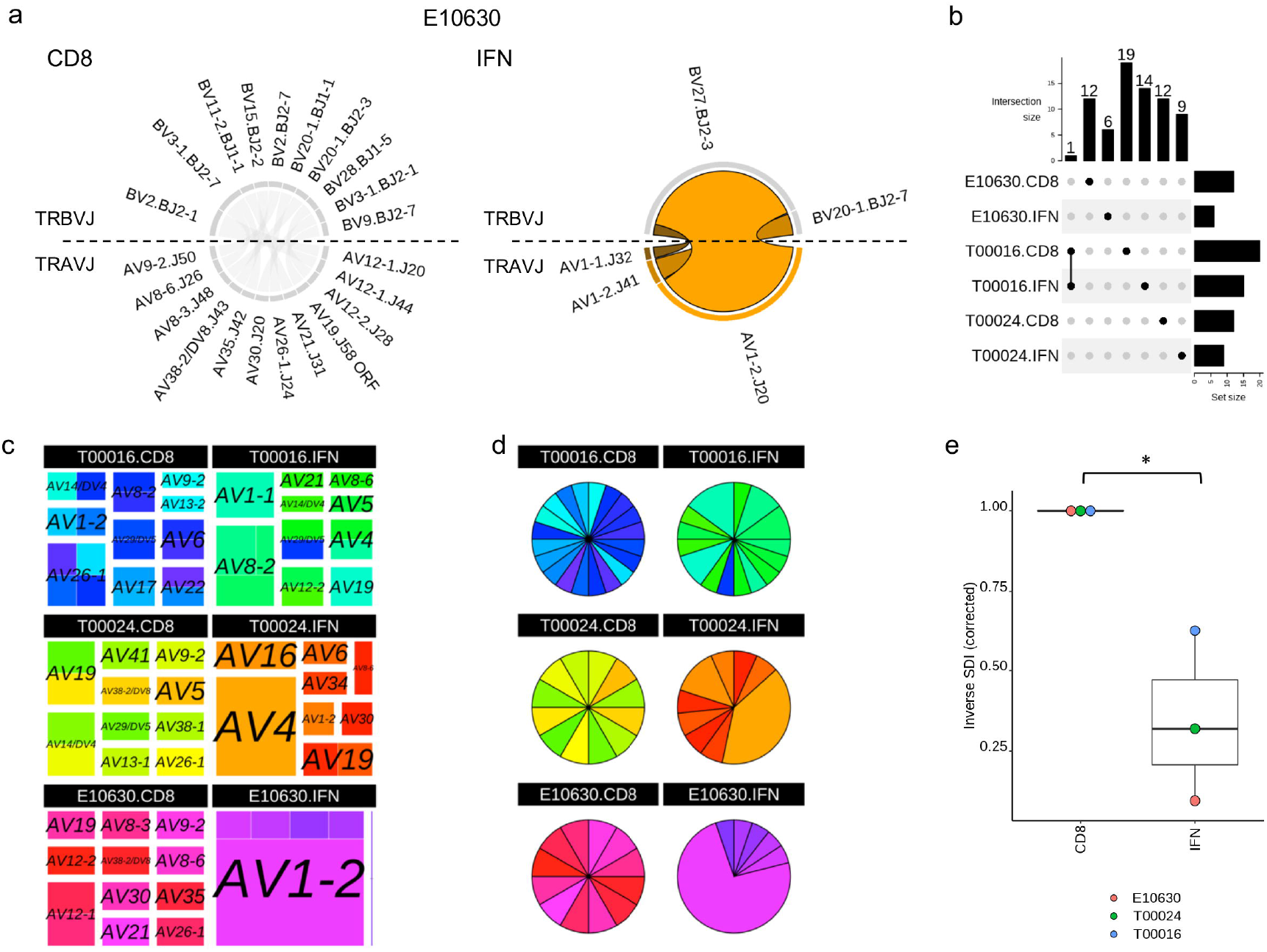
TCR-Explore analysis of a pre-existing TCR repertoire dataset. TCR repertoire data derived from *in vitro* expanded carbamazepine-induced T cells derived from patients with Stevens-Johnson Syndrome (T00016, T00024 and E10630), with CD8 and IFN representing the non-activated and drug-activated subsets, respectively. (**a**) E10630, chord diagram of drug-induced αβTCR repertoire for CD8 and IFN subsets. No overlapping sequences between the CD8 (grey) and IFN (orange) subsets. (**b**) Upset plot representing αβTCR CDR3 region overlap. Dots represent the presence of a clonal sequence and lines connect overlapping samples. (**c**) treemap coloured by AVJ_aCDR3_BVJ_bCDR3 and separated by the TRAV genes (size of the square indicate proportion of each TCR relative to the individual sample; colour represents a unique clone). (**d**) pie chart coloured by AVJ_aCDR3_BVJ_bCDR3 (size of the segment is proportional to the percentage of each clone; colour represents a unique clone). Same colours were used for both the (**c**) treemap and (**d**) pie chart. (**e**) Inverse Simpson index vs condition to measure change in diversity following drug exposure. Paired Students t-test, *p<0.05. Dots represent each individual.

There are three proposed mechanisms for T cell activation by small molecules drugs, with both the hapten/prohapten and altered repertoire mechanisms resulting in alteration to peptides presented by the HLA[15]. In contrast, our paper[14] showed that carbamazepine-induced SJS/TEN, associated with the pharmacological interaction with immune receptors concept, had minimal impact on the peptide repertoire. Therefore, T cell stimulation is likely to be triggered by direct interactions between the drug and the TCR, bypassing the need for a specific peptide/HLA complex. Using TCR-Explore, interrogation of CDR3α motifs (*i*.*e*. unique sequences) highlighted a redistribution of carbamazepine-induced CDR3α lengths towards 11mers (E10630) and 15mers (E10630, T00016), as well as loss of 16 mers (T00024) and 17mers (T00016, T00024) (**Figure 3a**). Interestingly, the E10630-derived 11mer (CAAFG**<u>D</u>**<u>YKLSF)</u> in our original publication was shown to be activated in the presence of both CBZ and HLA-B15:02. For the 15mers, the IFN activated group was dissimilar from the CD8 non-activated group for both T00016 (**Figure 3b**) and T00024 (**Figure 3c**; major clonotype CDR3α TRAV4-TRAJ33 CLVGETG**<u>D</u>**<u>SNYQLIW</u>). Interestingly, the E10630 and T00024 CDR3α two clonotypes shown above presented a central aspartate residue (bolded D). A centric aspartic acid residue αCDR3 region was also observed in carbamazepine-induced TCRs of other study participants (E10056, CAAKDG<u>M</u>**<u>D</u>**<u>SSYKLIF;</u> AP026, CIVRSLR**<u>D</u>**<u>NYGQNFVF</u>)[14] which were also activated in the presence of CBZ and HLA-B*15:02. Moreover, centric aspartic acid residues have been previously reported in carbamazepine-induced SJS/TEN blister fluid-derived T cells (VF**D**<u>NT</u>**<u>D</u>**<u>KLI</u> and AASPP**D**<u>GNQFY</u>)[59]. Therefore, the centric aspartic acid was most commonly derived from different TRAJ genes (underlined section), and this feature may be required for CBZ-induced TCR activation. Together, TCR_Explore provided an opportunity to further interrogate our dataset that contributed to novel insights and opened further avenues for functional investigation.

**Figure 3.**
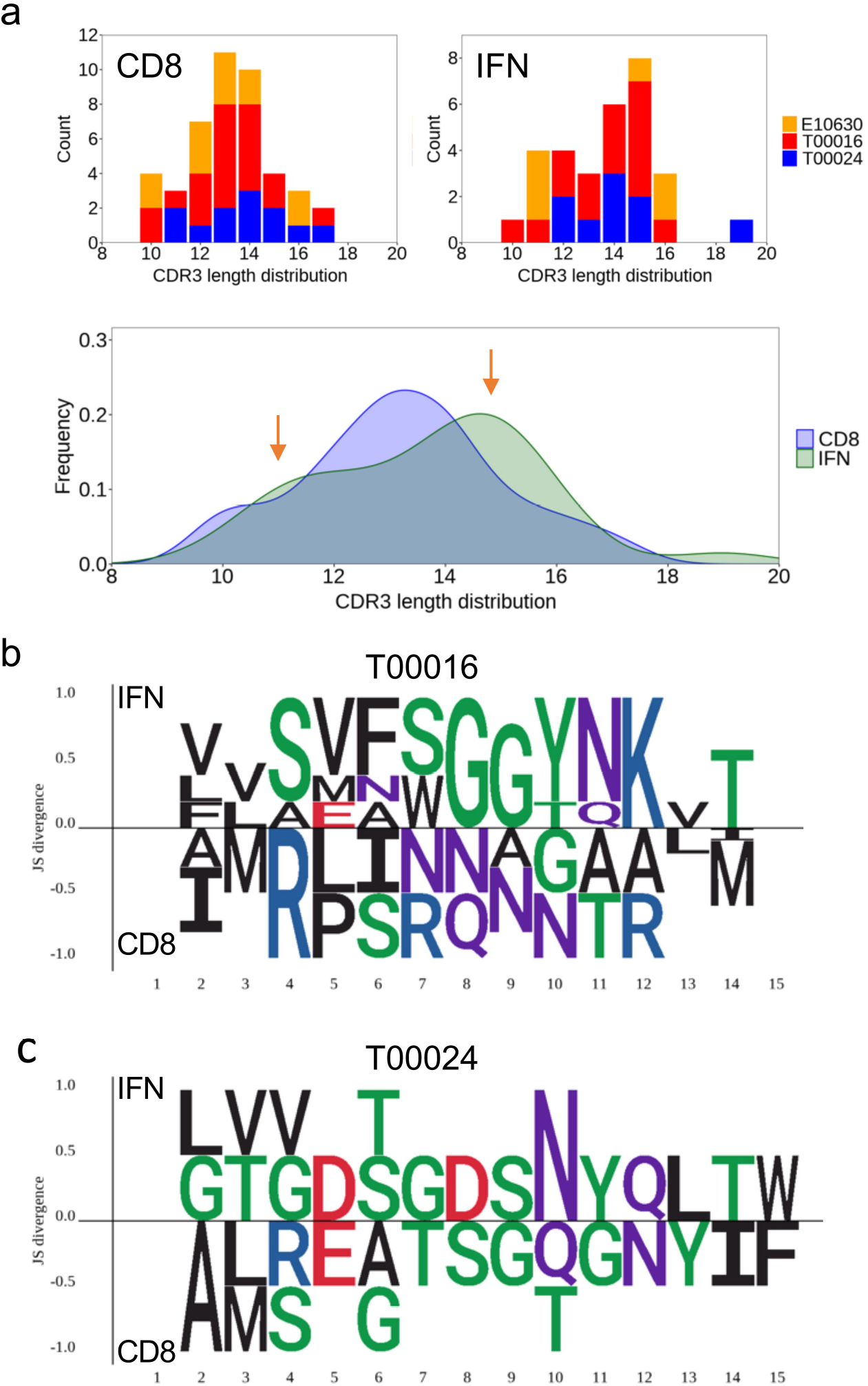
TCR-Explore reveals CDR3α motif nuances. TCR repertoire data from patients with Stevens-Johnson Syndrome (T00016, T00024 and E10630), with CD8 and IFN representing the non-activated and drug-activated subsets, respectively. (**a**) CDR3α length distribution coloured by individual or density plot of CD8 non-activated vs IFN activated group. CDR3α motif plot showcasing the IFN activate (top) vs CD8 non-activated (bottom) 15mer for (**b**) T00016 and (**c**) T00024.

### Linking of TCR clonotypes with immunophenotypes in a mouse model of autoimmunity

In another recent study[12], we examined the αβTCR repertoire in a mouse model of Rheumatoid Arthritis that expresses the human susceptibility allele HLA-DRB1*04:01. HLA-DR4 mice were inoculated with a double citrullinated peptide Fibβ-72,74cit_69-81_, and lymphocytes from draining lymph nodes were collected on day 8 and examined for TCR cross-reactivity to the single and double citrullinated epitopes by co-staining with Fibβ- 72,74cit_69-81_ and Fibβ-74cit_69-81_ tetramers. Individual unique- and cross-reactive CD4^+^ T cells were index sorted for downstream association of immunophenotype and TCR sequence and single cell TCRα and TCRβ sequencing. Concordant with the original analysis, TCR_Explore highlighted the cross-reactive αβTCR clone (*i*.*e*. TRAV1/J37-TRBV13-1/J1-6; green cross), as depicted by high tetramer expression of both single- and dual-stained citrullinated peptides (**Figure 4a**). However, additional information was showcased by TCR_Explore following further examination of the phenotyping panel (*e*.*g*. CD4, CD8, TCRβ, CD62L). Here, we confirmed that the immunophenotype of the tetramer sorted T cells were all CD4^+^ TCRβ^+^ (**Figure 4b**) and CD62L^low^ (**Figure 4c**). Low expression of CD62L has also been associated with T cell activation in rheumatoid arthritis[61]. Overall, TCR_Explore provides a critical platform to examine TCR signatures with immunophenotyping captured via FACSort index data to identify phenotypic markers of interest.

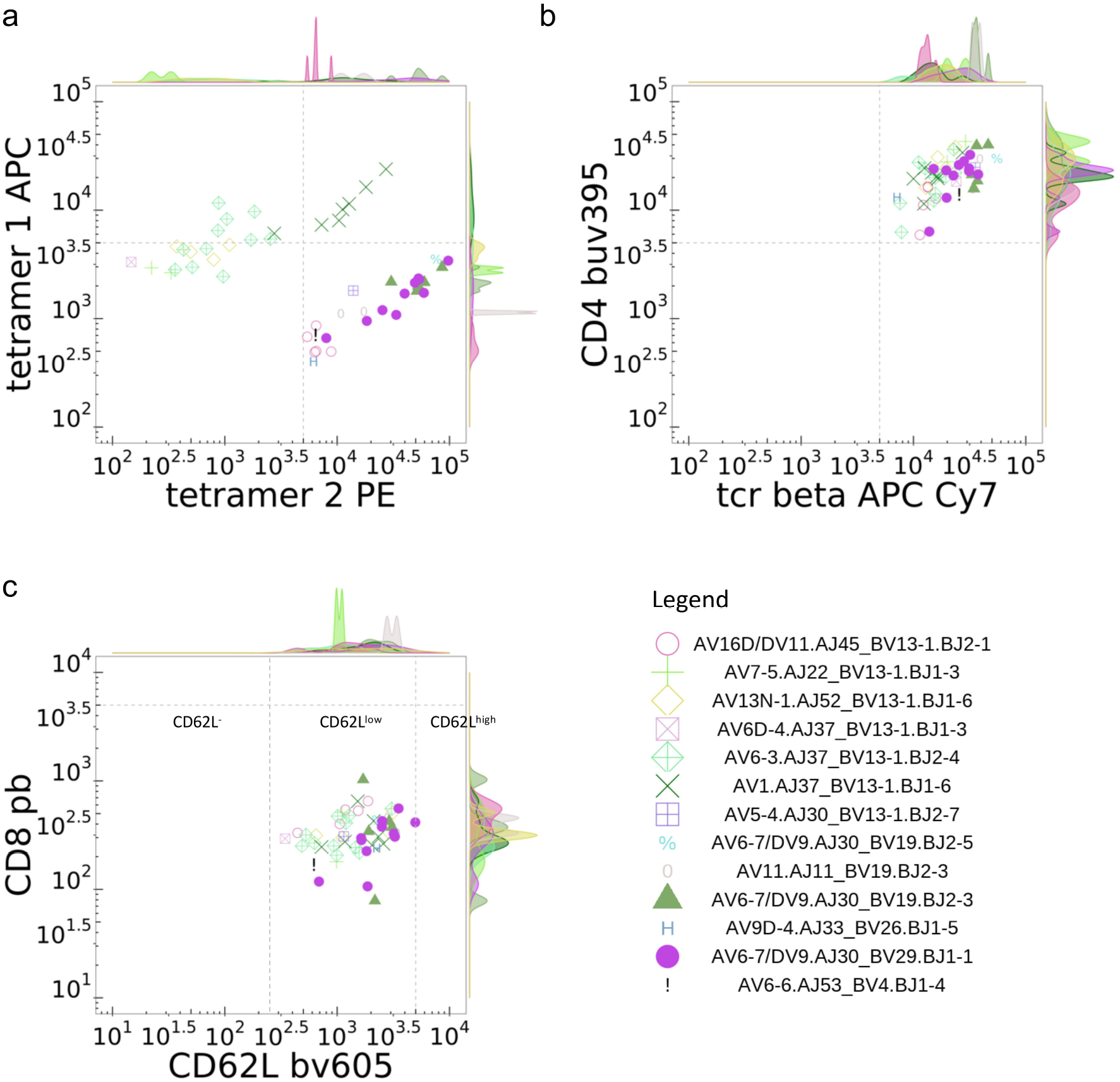

## Discussion

Programs that perform in-depth TCR repertoire analysis (*e*.*g*. TCRdist[19], TCRdist3[24], clusTCR[25], VDJtools[20]) utilise coding languages such as python and R, which effectively limits their usage to individuals with experience in these programming languages or at the very least requires a dedicated time commitment to learn these languages for accessibility. Alternatively, other programs such as Immunarch[23], a coding-based R tool, is aimed to improve access to TCR analysis by minimising the amount of coding needed to a maximum of 5-10 lines[23]. We have further improved the user experience of TCR repertoire analysis by launching our R application TCR_Explore on a website, which does not require coding inputs for program operation.

One of the most critical processes for any dataset analysis is QC of the raw data. Prior to TCR_Explore, our workflow involved the processing of Sanger sequencing information into IMGT to generate the TCR assignments via a vquest.xls file. This file then underwent manual QC to generate a curated file for TCR analysis, which increases the potential for data transcription errors to occur. To eliminate both manual QC processing and data transcription errors, TCR_Explore was purposefully designed to include an automated QC function, with greatly reduced errors and efficiency that enables users to analyse their data in a shorter timeframe.

Flexibility of data selection is an essential design feature in TCR_Explore to ensure that different variables can be examined without the need to modify the single curated input data file. Previous analyses that required changes to the dataset, such as the addition of individuals and/or treatment groups, would magnify the time required for reanalysis as this would also involve manual reformatting steps required for the various programs being used. TCR_Explore automates reformatting and eliminates the need for the modification of multiple files, enabling the user to interrogate their dataset more thoroughly in the first instance.

The capacity to visualise TCR repertoire data has often been restricted to programs and webtools that employ either coding languages and/or require specific file formats for input data, which also introduces the possibility of data transcription errors. Indeed, in some instances more than one webtool is required to visualise different graphical representations of the dataset. Here, TCR_Explore consolidates the generation of 15 different types of figures, with 14 of these able to be created from the same input file in the ‘TCR analysis’ section. To demonstrate the strength of this feature we showcased an increased flexibility and capacity to perform in-depth TCR repertoire analysis by re-examining our previously published human drug-induced TCR dataset[14]. Not only were we able to recapitulate our initial findings but extended our analysis in terms of evaluating both altered diversity of the TCR sequences and CDR3α motif differences. Access to all these figures in the one location enabled us to further interrogate our data with new hypotheses, which led to novel lines of inquiry not previously appreciated.

The capacity to pair the immunophenotype and TCR signature of an antigen-specific T cell provides powerful information for identification of disease biomarkers[58]. TCR_Explore was tailored to readily merge single-cell TCR sequencing and FACS Index sorted information. Re-evaluation of our mouse autoimmune TCR dataset[12] with TCR_Explore improved the robustness of data interrogation and visualisation to showcase previously unappreciated immunophenotypic markers of interest associated with rheumatoid arthritis. Our program facilitated the examination of two immunophenotypic biomarkers at a time, where the comparisons could be readily changed. Importantly, third party programs involving paid subscriptions or coding-based programs are no longer required to perform this function. Our program has improved capacity to identify T cell biomarkers that could be used in disease diagnoses, as well as identification of immunogenic T cells that have the potential to be developed into T cell-based therapeutics.

## Conclusion

TCR_Explore is a purpose-designed program to perform automated TCR repertoire analysis and visualisation. TCR_Explore includes a QC pipeline to aid in error-free and proficient TCR repertoire analysis, as well as the generation of a single input file for data analysis and creation of publication ready figures. Use of the Shiny R interface and program maintenance on a webserver ensures that TCR_Explore is accessible to users irrespective of their coding expertise. We anticipate that TCR_Explore will provide a powerful platform for interrogation of TCR repertoires to unravel the complexity of their contribution in human health and disease.

## Acknowledgements

**KAM** would also like to acknowledge her additional PhD supervisors Prof. Patrick Kwan and Dr Alison Anderson for their support during her candidature.

## Author Contribution

**KAM** contributed to the Conceptualization, Methodology, Software, Validation, Writing original draft. **NAM** contributed to the Conceptualization, Validation, Writing review & editing. **CL** contributed to the Resources, Validation, Writing review & editing. **JR and AWP** contributed to the Resources, Writing review & editing. **CJ, JBZ, SJRG, PTI** and **NLG** contributed to Validation, Writing review & editing.

## Ethics declaration

The authors declare that they have no known competing financial interests or personal relationships that could have appeared to influence the work reported in this paper.

## Supplementary information

Three supplementary data files and three supplementary tables.

